# (-)-Gossypol inhibition of musashi-mediated forgetting improves memory and age-dependent memory decline in *Caenorhabditis elegans*

**DOI:** 10.1101/2022.06.20.496809

**Authors:** Pavlina Mastrandreas, Andreas Arnold, Csaba Boglari, Dominique J.-F. de Quervain, Attila Stetak, Andreas Papassotiropoulos

## Abstract

Musashi RNA-binding proteins retain a pivotal role in stem cell maintenance, tumorigenesis, and nervous system development. Recently, we showed in *C. elegans* that MSI1 actively promotes forgetting upon associative learning via a 3’UTR-dependent translational expression of the Arp2/3 actin branching complex. Here, we investigated the evolutionary conserved role of MSI proteins and the effect of their pharmacological inhibition on memory. Expression of human MSI1 and MSI2 under the endogenous musashi promoter fully rescued the phenotype of *msi-1(lf)* worms. Furthermore, pharmacological inhibition of MSI1 and MSI2 activity using (−)-gossypol resulted in improved memory retention, without causing locomotor, chemotactic, or learning deficits. No drug effect was observed in *msi-1(lf)* treated worms. Using Western blotting and confocal microscopy we found no changes in MSI-1 protein abundance following (−)-gossypol treatment, suggesting that musashi gene expression remains unaltered and that the compound exerts its inhibitory effect post-translationally. Additionally, (−)-gossypol suppressed the previously seen rescue of the *msi-1(lf)* phenotype in worms expressing human MSI1 specifically in the AVA neuron, indicating that (−)-gossypol can regulate the musashi pathway in a memory-related neuronal circuit in worms. Finally, treating aged worms with (−)-gossypol reversed physiological age-dependent memory decline. Taken together, our findings indicate that pharmacological inhibition of musashi might represent a promising approach for memory modulation.

## Introduction

Learning and memory constitute cognitive functions comprising of a wide variety of complex components. Even though there has been a lot of effort to elucidate the molecular aspects of memory (Squire, 1987; Wang and Morris, 2010; Kandel et al., 2014), actively controlled forgetting is an equally important but much less investigated mechanism (Kraemer and Golding, 1997; Davis and Zhong, 2017; Ryan and Frankland, 2022).

We have recently demonstrated in *C. elegans* that the musashi RNA-binding protein (RBP) plays an important regulatory role in forgetting by modulating cytoskeletal changes at the translational level (Hadziselimovic et al., 2014). The musashi protein was originally discovered as a key player in asymmetric cell division, stem cell function, and cell fate determination in *Drosophila* (Okano et al., 2005). In vertebrates, the two *MSI* orthologues, *musashi1* (*MSI1*) and *musashi2* (*MSI2*) are mostly expressed in stem cells. In particular, MSI1 is a marker of neural progenitor cells (Sakakibara et al., 1996), and outside the nervous system can be found in the stem cells of the gut (Kayahara et al., 2013) and epithelial cells of the mammary gland (Colitti and Farinacci, 2009). MSI2 is also present in differentiated neurons of the adult brain as well as in hematopoietic stem cells (Kharas et al., 2010). Expression of MSI1 has been correlated with the grade of the malignancy and proliferative activity in gliomas and melanomas, whilst mutations in MSI2 have been associated with poor prognosis in certain types of cancers. Both MSI1 and MSI2 harbor two tandem RNA recognition motifs (RRMs) located at their N-terminal region, followed by a putative disordered region. The amino acid sequences of RRM1 and RRM2 of MSI1 exhibit 85% identity to those of MSI2 respectively, suggesting that the two proteins might target some mRNAs in a similar fashion. Indeed, an *in vitro* study identified the UAG RNA sequence as the core MSI1 binding motif (Zearfoss, 2014), while recent genome-wide analyses of MSI2 targets revealed binding of MSI2 to mRNA 3’UTRs that contain multiple copies of UAG motifs as well as polyU pentamers (Bennett, 2016; Karmakar et al., 2022).

Dysregulated expression of RBPs can be detected in various human diseases ranging from neurological disorders to cancer (Lukong et al., 2008; Darnell, 2010), rendering them potential targets for therapeutic intervention. However, identifying compounds that can disrupt protein-RNA interactions remains an arduous task due to the complexity of targeting RBPs or in the selection of their interacting mRNAs. Previous studies have shed light on small molecule inhibitors of MSI1 in an effort to reduce musashi-mediated tumourigenesis. Ω-9 fatty acids, such as oleic acid and its derivatives, were identified as allosteric inhibitors of MSI1 RNA binding activity (Clingman et al., 2014), while a fluorescence polarization (FP) competition assay revealed that (−)- gossypol, a natural compound extracted from cottonseed, acts as an *in vitro* inhibitor of MSI1 by recognizing and directly binding to its RNA binding domain 1 (RBD1) (Lan et al., 2015).

*Caenorhabditis elegans* represents a powerful model organism to elucidate the molecular mechanisms of learning and memory. It is characterized by a simple nervous system of 302 neurons capable of exhibiting behavioral plasticity towards an array of stimuli (Colbert and Bargmann, 1995; Sengupta et al., 1996). In *C. elegans,* the sole musashi ortholog *(msi-1)* has been implicated in male mating (Yoda et al. 2000). We previously demonstrated that MSI-1 actively regulates forgetting via translational repression of actin branching complexes (Hadziselimovic et al., 2014). (−)- gossypol was the first small molecule compound to exhibit potent anticancer activity *in vivo* by disrupting MSI1 activity via the Notch/Wnt pathway (Lan et al., 2015), rendering it an ideal candidate to test for its potential effects on memory. In the current study, we investigated the evolutionary conserved role of MSI proteins and the effect of their pharmacological inhibition on memory and physiological age-dependent memory decline in these animals.

## Results

### Human MSI1 and MSI2 proteins are implicated in memory

Previously, we have shown that the sole *C. elegans* musashi ortholog, *msi-1*, actively promotes forgetting (Hadziselimovic et al., 2014). To study the potentially conserved role of MSIs in memory we performed rescue experiments with the human musashi homologues MSI1 and MSI2 in *C. elegans,* using established context-dependent associative learning and memory assays (Nuttley et al. 2002, Vukojevic et al., 2012).

In order to study the functional conservation of musashi proteins between worms and humans (Figure 1A), we generated transgenic *C. elegans* lines expressing human MSI1 and MSI2 in a *msi-1* loss of function *(lf)* background. The transgenic animals carried a 7.7 kb *C. elegans msi-1* promoter region fused to the human *MSI1* or *MSI2* cDNA followed by 1.1 kb of the *C. elegans msi-1* 3’UTR. Following the integration of the arrays, the expression of both transgenes was confirmed with RT-PCR (Figure 1B), and the lines were tested in a short- and long-term negative olfactory learning and memory paradigm. Initially, wild-type, *msi-1(lf)*, *msi-1(lf); Is[hsmi1+]* and *msi-1(lf); Is[hsmi2+]* animals were tested for their chemotaxis toward diacetyl (DA) prior to and following aversive associative learning (conditioning). All animals exhibited similar chemotaxis, independent of the genotype tested (Figure 2A-D), suggesting that ectopic expression of human MSIs in worms does not influence baseline chemotaxis or associative learning. Moreover, we investigated the role of human MSI1 and MSI2 in short-term [STAM] and long-term associative memory [LTAM]. In the case of STAM, worms were subjected to one round of conditioning and tested for memory performance following a 1h recovery period (Nuttley et al., 2002). As previously shown, *msi-1(lf)* exhibited increased memory retention in short-term memory (Hadziselimovic et al., 2014), a phenotype that was fully restored to wild-type levels in *msi-1(lf)* mutants carrying either the human *MSI1* or *MSI2* transgene (Figure 2A, B). Similar to short-term memory, expression of either human *MSI*s fully rescued the effect of *msi-1(lf)* mutant worms on memory retention after a 24h delay period (Figure 2C, D) (Vukojevic et al., 2012, Hadziselimovic et al., 2014). Altogether, our results indicate that both human musashi homologues can replace *C.elegans* MSI-1 protein function in STAM as well as in LTAM.

**Figure 1.**
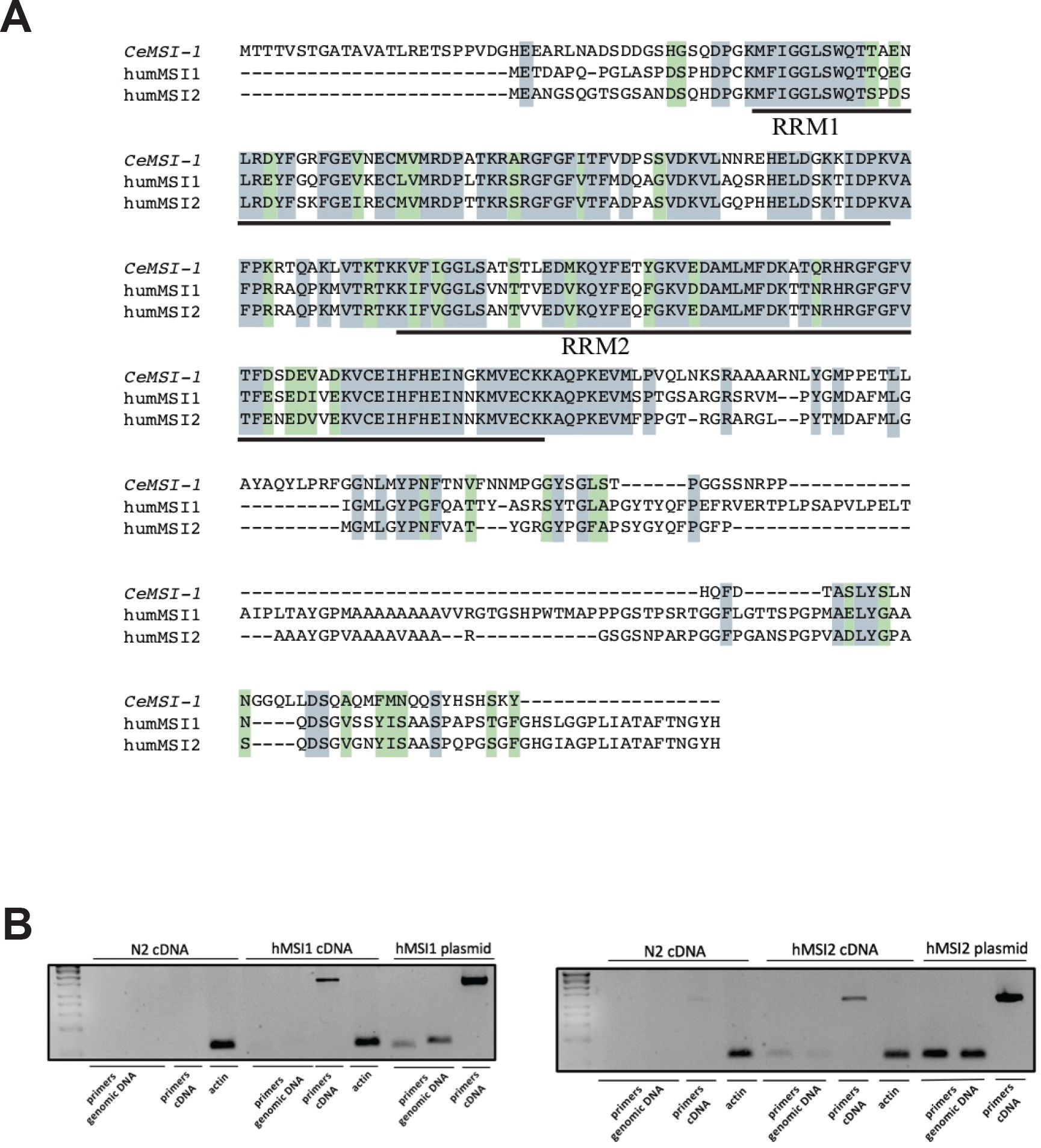
Sequence alignment of *C. elegans* and human musashi orthologues and validation of transgene expression. A. Grey boxes indicate identical amino acids and green boxes show similar amino acids. The two RNA-binding domains (RRM1 and RRM2), showing the highest conservation amongst species, are underlined. B. Expression of human MSI1 and MSI2 was validated using cDNA extracted from the transgenic lines. N2 cDNA was used as a negative control for musashi expression and the original plasmids used for cloning were used as positive controls for primer annealing. Actin was used as a further control for cDNA integrity. Primers annealing to the endogenous *msi-1* promoter/ human cDNA boundary identify any amplification coming from contaminating genomic DNA (“primers genomic DNA”) and primers binding within the human cDNA region illustrate true expression (“primers cDNA”).

**Figure 2.**
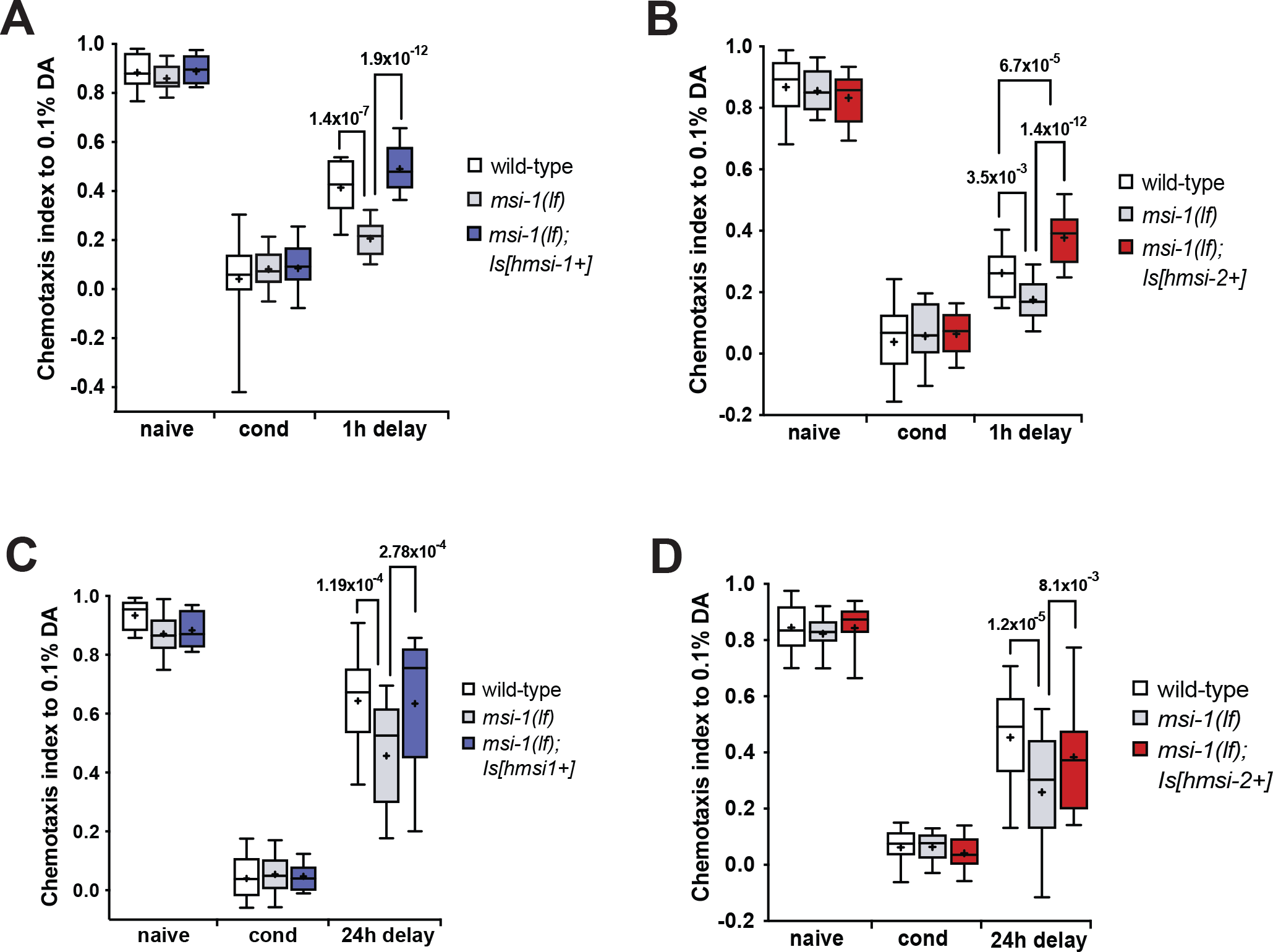
Human MSI1 and MSI2 are involved in the regulation of memory loss. (A-B) Negative STAM was tested in wild-type, *msi-1(lf),* and *msi-1(lf)* mutant worms rescued with the human *MSI1* or *MSI2* construct. Worms were assayed toward DA without (naïve), with preincubation with DA and starvation (cond) or after 1h (1h delay). (C-D) Negative LTAM in the different genotypes was tested following two consecutive conditioning phases and DA preference was tested immediately (cond) and after a 24h (24h delay) recovery period. All experiments were done in triplicates and repeated at least five times. Box plots for each genotype and condition are presented with whiskers indicating the 10th and 90th percentiles. Statistical significance between groups was assessed with two-way ANOVA and post hoc t-tests as indicated (Bonferroni’s adjusted p values are reported).

### Human MSI1 protein is necessary in the AVA interneuron

Previously, it was shown that expression of the *C. elegans* MSI-1 under the *rig-3* promoter, which drives expression primarily in the AVA interneuron, was sufficient to rescue the memory phenotype of *msi-1(lf)* mutants (Hadziselimovic et al., 2014). To find out whether the same was true for human musashi, we performed a tissue-specific rescue experiment by expressing the human *MSI1* cDNA under the control of the *C. elegans rig-3* promoter in *msi-1(lf)* worms. We could observe rescue in both the STAM (Figure 3A) as well as in the LTAM (Figure 3B) testing paradigm.

**Figure 3.**
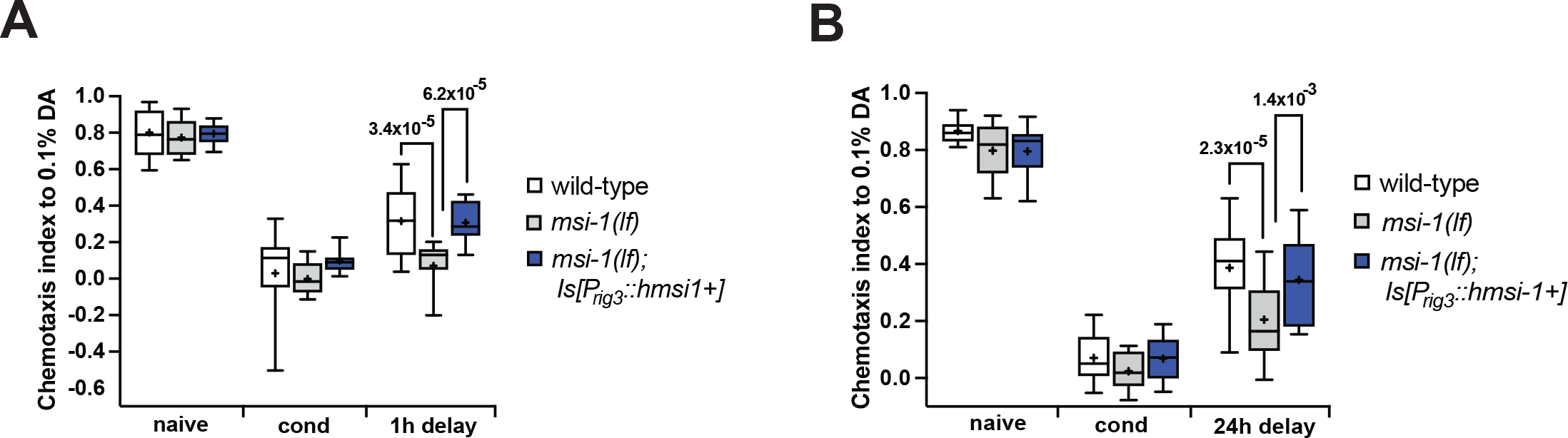
Human MSI1 regulates memory loss in the AVA interneuron. (A) Negative STAM was tested in wild-type, *msi-1(lf),* and *msi-1(lf)* mutant worms expressing the human *MSI1* construct. Worms were assayed toward DA without (naïve), with preincubation with DA and starvation (cond) or after 1h (1h delay). (B) Negative LTAM in the different genotypes was tested following two consecutive conditioning phases and DA preference was tested immediately (cond) and after a 24h (24h delay) recovery period. All experiments were done in triplicates and repeated at least five times. Box plots for each genotype and condition are presented with whiskers indicating the 10th and 90th percentiles. Statistical significance between groups was assessed with two-way ANOVA and post hoc t-tests as indicated (Bonferroni’s adjusted p values are reported).

### (−)- gossypol has no adverse effect on the chemotactic or locomotory behavior of C. elegans

Surface plasmon and nuclear magnetic resonance studies have demonstrated that (−)- gossypol directly interacts with the RNA binding groove of MSI1 and disrupts binding to its downstream targets (Lan et al., 2015). (−)- gossypol’s therapeutic applicability has been limited due to the toxicity it harbors (Gadelha et al., 2014). In order to address any potential toxic effects, we first measured the chemotactic response of worms towards diacetyl following treatment with different gossypol concentrations. In both human *MSI1* and *MSI2* transgenic worms, we observed no significant immediate (Figure 4A, B) or delayed changes (Figure 4C, D) in the chemotaxis index of worms treated with (−)- gossypol compared to untreated DMSO controls.

**Figure 4.**
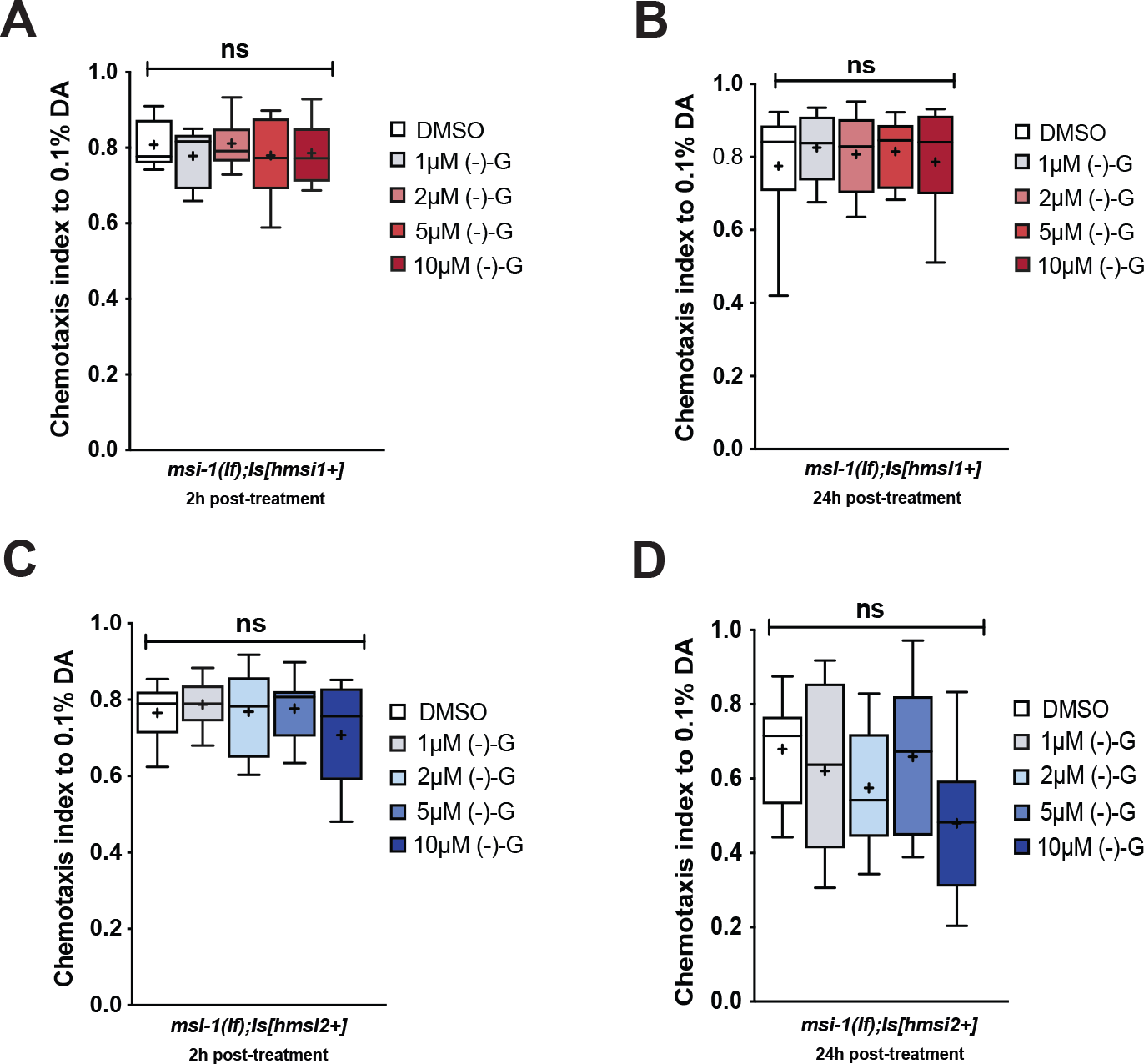
(-)- gossypol harbors no immediate or delayed toxicity. (A-B) Naïve chemotaxis was tested in *msi-1(lf)* mutant worms rescued with the human MSI1 construct following a 2h (A) and 24h (B) delay, post-(-)-gossypol treatment. (C-D) Naïve chemotaxis was tested in *msi-1(lf)* mutant worms rescued with the human MSI2 construct following a 2h (C) and 24h (D) delay post- (-)-gossypol treatment. All experiments were done in triplicates and repeated at least five times. Box plots for each genotype and condition are presented with whiskers indicating the 10th and 90th percentiles. Statistical significance between groups was assessed with two-way ANOVA and post hoc t-tests as indicated (Bonferroni’s adjusted p values are reported) ns: not significant.

To determine whether (-)- gossypol modulates the locomotory behavior of *C. elegans,* we measured the number of body bends per minute in worms treated with DMSO as a vehicle control and 10μM (-)- gossypol. Following a two-hour drug treatment, worms were transferred to clean OP50 seeded plates and allowed to move freely for three minutes before counting the locomotory rate. Using this measure, we found no difference in the locomotor response of *C. elegans* exposed to 10μM (-)- gossypol versus the control group (Figure 5A).

**Figure 5.**
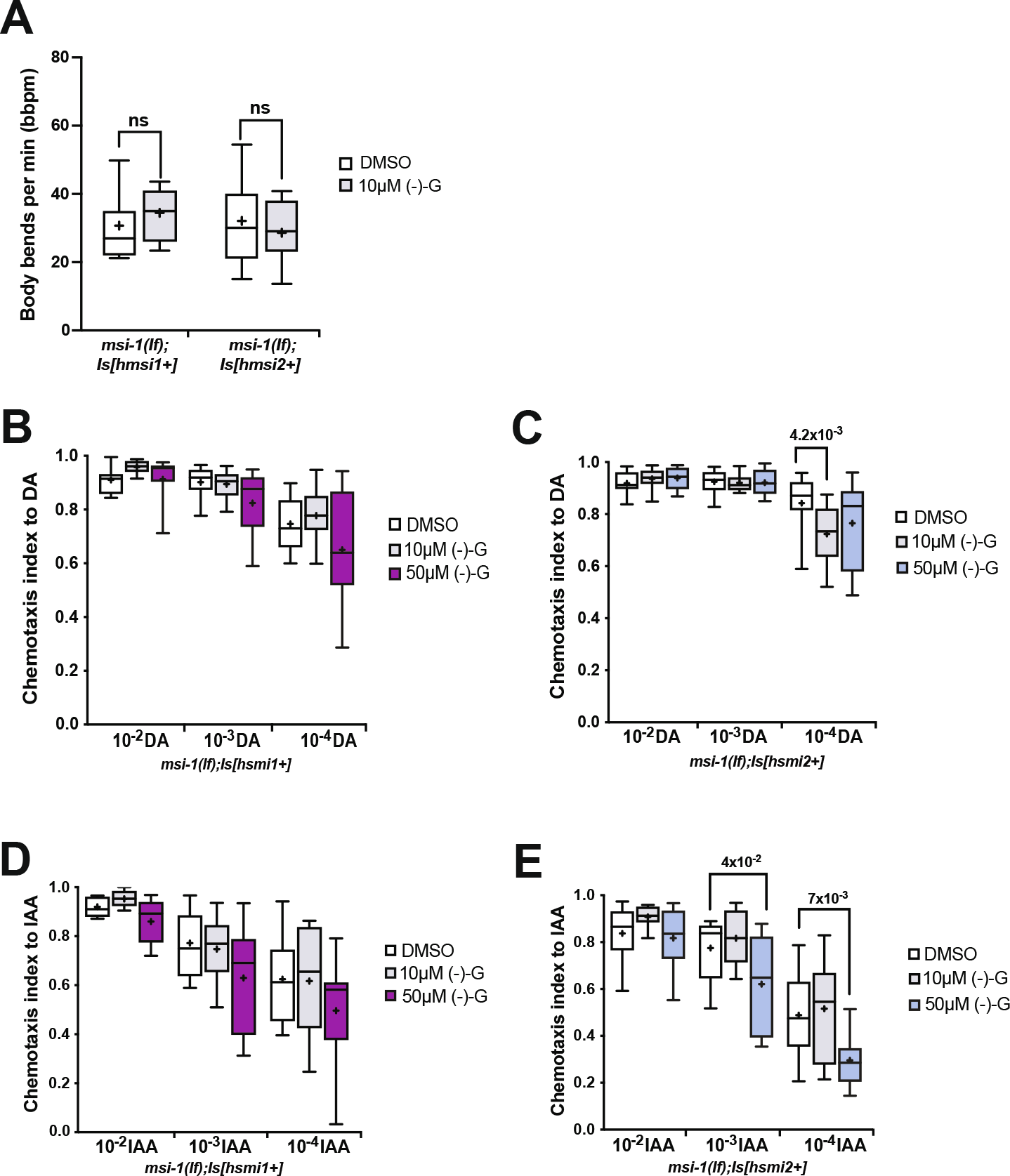
Effects of (-)- gossypol on the motoric and chemosensation behavior of worms. (A) Locomotory rate of DMSO and (-)- gossypol treated *msi-1(lf)* worms rescued with human MSI1 and human MSI2. Data were analyzed using a two-tailed Student’s test with p <0.05. Box plots for each genotype and condition are presented with whiskers indicating the 10th and 90th percentiles. (B-C) Chemotaxis after (-)- gossypol treatment was measured towards DA as previously described (Bargmann et al., 1993) in *msi-1(lf)* worms rescued with (B) human MSI1 and (C) human MSI2. (D-E) Chemotaxis after (-)- gossypol treatment was measured towards IAA as previously described (Bargmann et al., 1993) in *msi-1(lf)* worms rescued with (D) human MSI1 and (E) human MSI2. All experiments were done in triplicates and repeated at least five times. Box plots for each genotype and condition are presented with whiskers indicating the 10th and 90th percentiles. Statistical significance between groups was assessed with two-way ANOVA and post hoc t-tests as indicated (Bonferroni’s adjusted p values are reported), ns: not significant.

To investigate whether (-)- gossypol influences the chemosensation of worms, we measured the chemotactic response of DMSO-, 10μM, and 50μM (-)- gossypol-treated worms towards different concentrations of diacetyl (DA) and isoamyl-alcohol (IAA). In *msi-1(lf)* worms rescued with the human *MSI1* construct, (-)- gossypol did not affect chemosensation at any given concentration of drug or chemoattractant (Figure 5B, D). In *msi-1(lf)* worms rescued with the human MSI2 construct, we observed a reduction in the chemotaxis towards the lowest DA concentration (10^−4^) tested when worms were treated with 10μM of (-)- gossypol (Figure 5C). In contrast to DA, a high concentration of (-)- gossypol (50μM) treatment significantly reduced the chemotactic response of *msi-1(os1);utrIs14[pmsi-1∷human msi2, psur-5∷dsred]* worms towards 10^−3^ and 10^−4^ IAA concentrations (Figure 5E), suggesting that higher (−)-gossypol concentrations can impair the ability of these worms to detect low concentrations of odors. It is important to note that DA and IAA are detected via the AWA and AWC amphid sensory neurons respectively (Bargmann et al., 1993); therefore, (-)- gossypol likely affects chemosensation independent of the sensory neuron implicated.

### (-)- gossypol inhibits musashi-mediated forgetting and improves short- and long-term memory

Even though it is widely known that (-)- gossypol exhibits potent anti-tumor activity *in vivo,* no research has been done to shed light on its potential effects on memory. Thus, in order to pharmacologically modify MSI activity and memory, we investigated the possible effect of (-)- gossypol treatment on the memory performance of human *MSI1-* and *MSI2-*expressing *msi-1(lf)* animals. Treating worms with (-)- gossypol for 2 hours prior to conditioning induced suppression of the phenotype observed in human MSI1-carrying worms compared to the DMSO treated line, thus improving memory in the STAM and LTAM behavioral assay (Figure 6A, B). Similar findings were observed in human MSI2 expressing *C. elegans* lines, where (-)- gossypol inhibited the previously seen rescue of the forgetting defect phenotype, both in a STAM and LTAM behavioral assay (Figure 6C, D). Thus, (-)- gossypol was able to improve both short and long-term memory in either of the humanized *C.elegans* strains.

**Figure 6.**
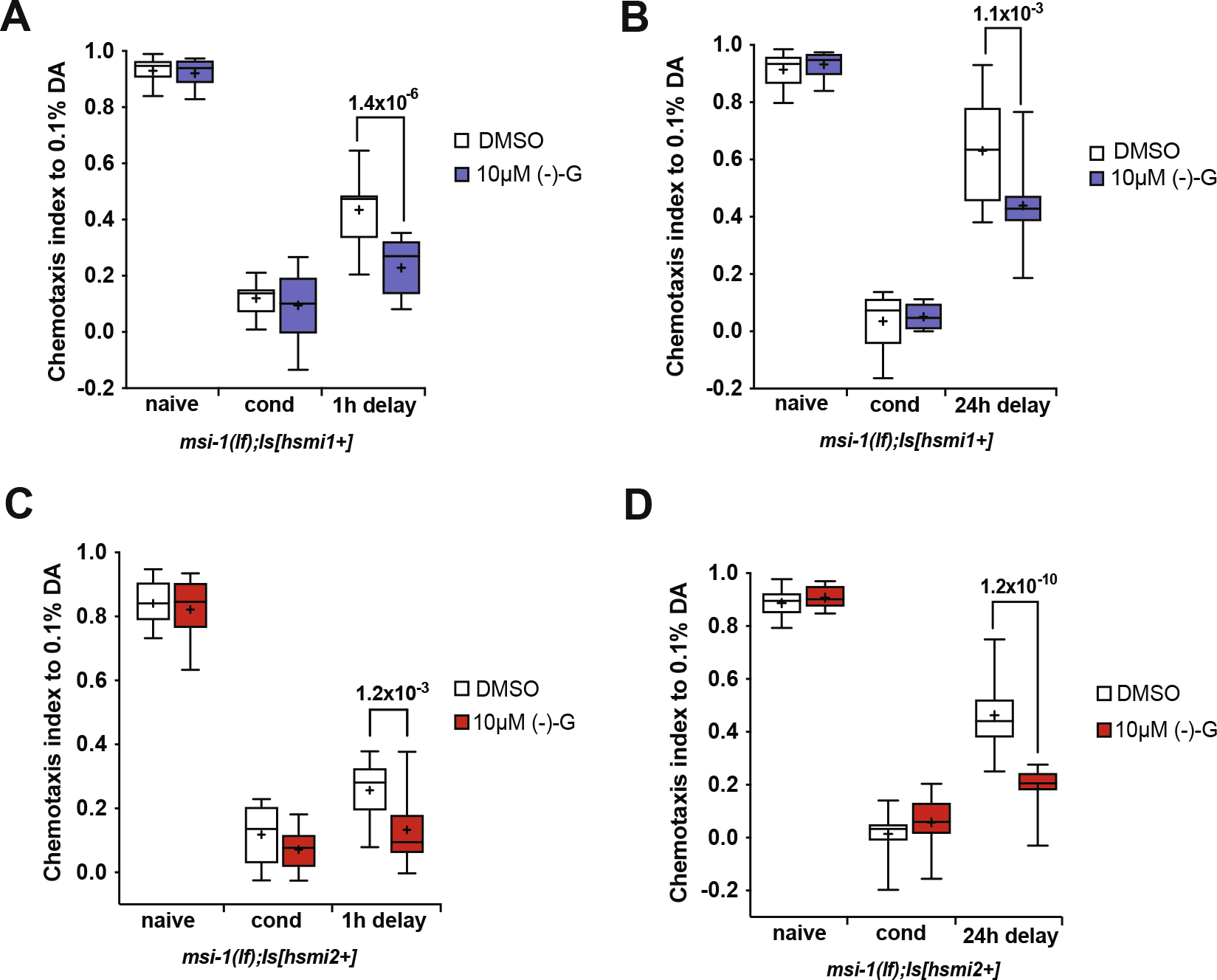
(-)- gossypol inhibits the memory rescue phenotype of human MSI1 and MSI2. (A-B) Negative STAM and LTAM were tested in *msi-1(lf)* mutant worms rescued with the human MSI1 construct or (C-D) with the human MSI2 construct; following a 2h (-)- gossypol treatment. Worms were assayed toward DA without (naïve) or with preincubation with DA and starvation (cond) and after 1h (1h delay). Negative LTAM in the different genotypes was tested following two consecutive conditioning phases and DA preference was tested immediately (cond) and after a 24h (24h delay) recovery period. All experiments were done in triplicates and repeated at least five times. Box plots for each genotype and condition are presented with whiskers indicating the 10th and 90th percentiles. Statistical significance between groups was assessed with two-way ANOVA and post hoc t-tests as indicated (Bonferroni’s adjusted p values are reported).

### (-)- gossypol-induced memory improvement is likely mediated through neuron-specific musashi inhibition

It was previously shown that (-)- gossypol does not exclusively inhibit MSI1 activity, but can also act on other proteins, such as on the BCL-2 family members (Oliver et al., 2004, Lian et al., 2012). To investigate the specificity of (-)- gossypol on MSI in the regulation of memory, we treated wild-type and *msi-1(lf)* worms with either DMSO or 10μM (-)- gossypol and measured the effect on memory after a 24h delay. While wild-type worms had a significant increase in LTAM retention when compared to their DMSO-treated control counterpart, *msi-1(lf)* worms displayed no differences between treatment groups (Figure 7A).

**Figure 7.**
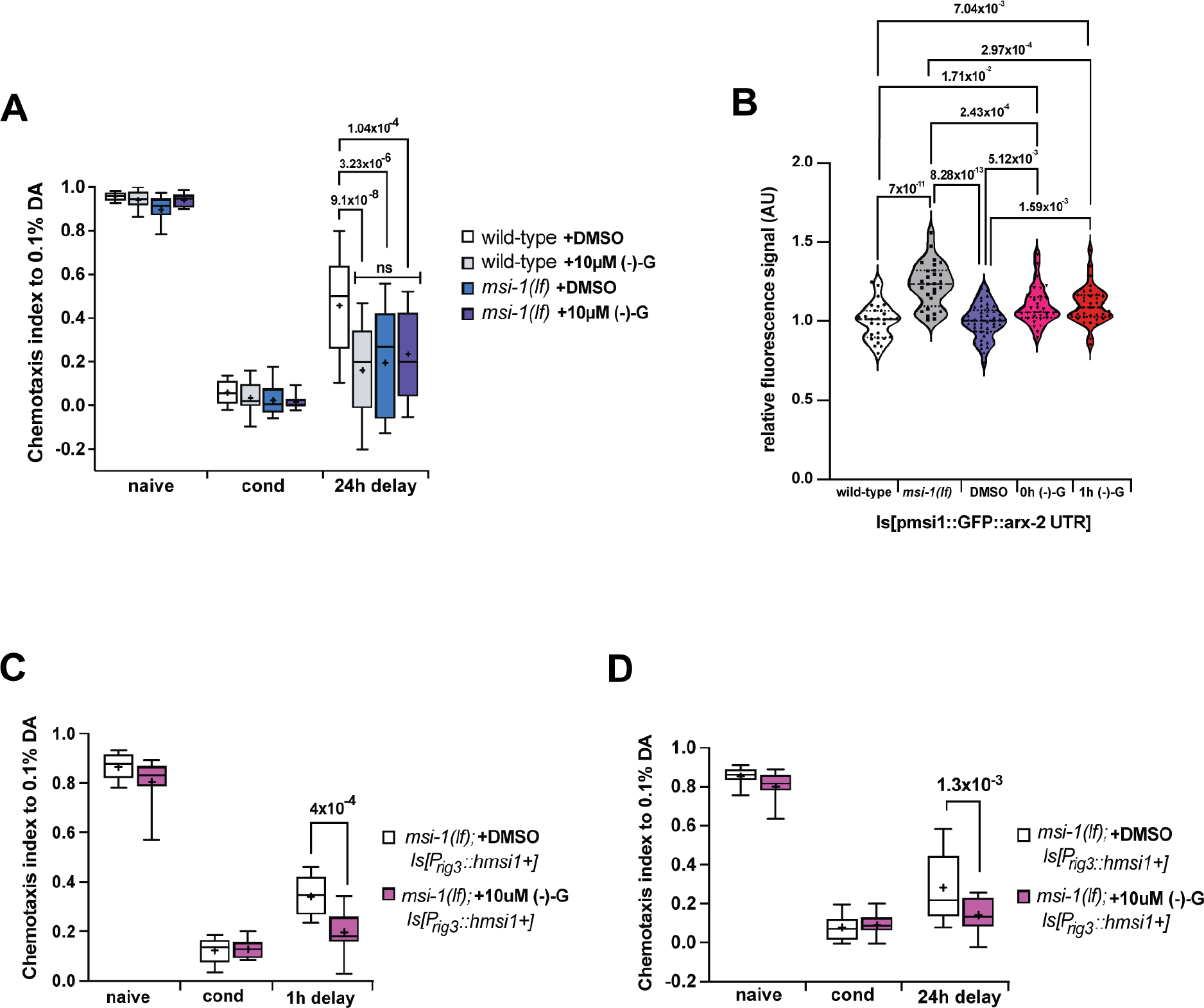
(-)- gossypol acts via the musashi pathway to inhibit forgetting. (A) Negative LTAM was tested in wild-type and *msi-1(lf)* worms treated with DMSO and 10μM (-)- gossypol. (B) GFP intensity in integrated transgenic worms carrying 7.7 kb *msi-1* promoter, GFP, and *arx-2* 3’ UTR. GFP signal was measured in untreated wild-type and *msi1 (lf)* worms and wild-type worms treated with either DMSO or (-)- gossypol (immediately or 1h hour after treatment). Box plots for each genotype and condition are presented with whiskers indicating the 10th and 90th percentiles. Statistical significance between groups was assessed with one-way ANOVA and post hoc t-tests as indicated (Bonferroni’s adjusted p values are reported). (C) Negative STAM was tested in *msi-1(lf)* mutant worms expressing human MSI1, treated with either DMSO or 10μM (-)- gossypol. Worms were assayed toward DA without (naïve) or with preincubation with DA and starvation (cond) and after 1h (1h delay). (D) Negative LTAM of treated worms was tested following two consecutive conditioning phases and DA preference was tested immediately (cond) and after a 24h (24h delay) recovery period. All experiments were done in triplicates and repeated at least five times. Box plots for each genotype and condition are presented with whiskers indicating the 10th and 90th percentiles. Statistical significance between groups was assessed with two-way ANOVA and post hoc t-tests as indicated (Bonferroni’s adjusted p values are reported).

Moreover, we investigated whether musashi downstream targets change upon (-)- gossypol treatment. We focused on changes of the Arp2/3 actin member ACTR2/ARX-2, previously reported to interact with MSI-1 (Hadziselimovic et al., 2014, Yang et al., 2019) and whose abundance significantly increased in *msi-1 (lf)*worms (Hadziselimovic et al., 2014) and upon (-)- gossypol treatment (Yang et al., 2019). We treated worms expressing the transgene pmsi-1∷GFP∷arx-2 3’UTR with either DMSO or 10μM (-)- gossypol for two hours and quantified GFP levels of head neurons immediately or 1h hour after treatment. We also measured GFP signals of the transgene in untreated wild-type and *msi-1 (lf)* worms as a control. In line with previous results, we observed a highly significant increase in the GFP signal in *msi-1 (lf)* worms compared to wild-type worms (Figure 7B). DMSO-treated worms showed similar GFP expression levels to those of wild-type worms suggesting that DMSO treatment alone does not affect ARX-2 protein levels (Figure 7B). Wild-type worms treated with 10μM (-)- gossypol showed a significant increase in GFP expression compared to untreated and DMSO-treated wild-type worms (Figure 7B), suggesting that a two-hour treatment with -(−) gossypol is sufficient to partially inhibit musashi and influence expression of its downstream targets.

The *msi-1* promoter has been suggested to drive expression in many tissues such as multiple neurons, muscle cells, and the intestine (wormbase.org) which can have implications for the drug response. Consequently, we investigated whether (-)- gossypol has a tissue-specific mode of action rather than an off-target effect. We treated *msi-1(lf); prig-3∷human msi1cDNA∷msi-1 3’UTR* worms with either DMSO or 10μM (-)- gossypol and measured the memory phenotype after a 1h or 24h delay. In both the STAM (Figure 7C) and LTAM (Figure 7D) paradigms tested, (-)- gossypol treatment was sufficient to suppress the *msi-1* rescue phenotype previously shown in the AVA interneuron, further validating that the effect of (-)- gossypol on memory is unlikely driven by alternative, off-target pathways.

### (-)- gossypol treatment does not influence MSI-1 protein abundance

To investigate whether any of the effects of (-)- gossypol could be a result of altered protein expression rather than a result of inhibition of protein function, we conducted a time course assay and examined MSI-1 endogenous protein levels after treatment. We first tagged the endogenous gene at the N-terminal end with a yellow fluorescent protein (YPET) and 3xFLAG using CRISPR/Cas9 (Dickinson et al., 2015). To confirm the functional integrity of the tagged protein, we compared the memory performance between wild-type, *msi-1(lf)* and MSI-1∷YPET∷3xFLAG animals and found no difference in either STAM or LTAM (Figure 8A, B).

**Figure 8.**
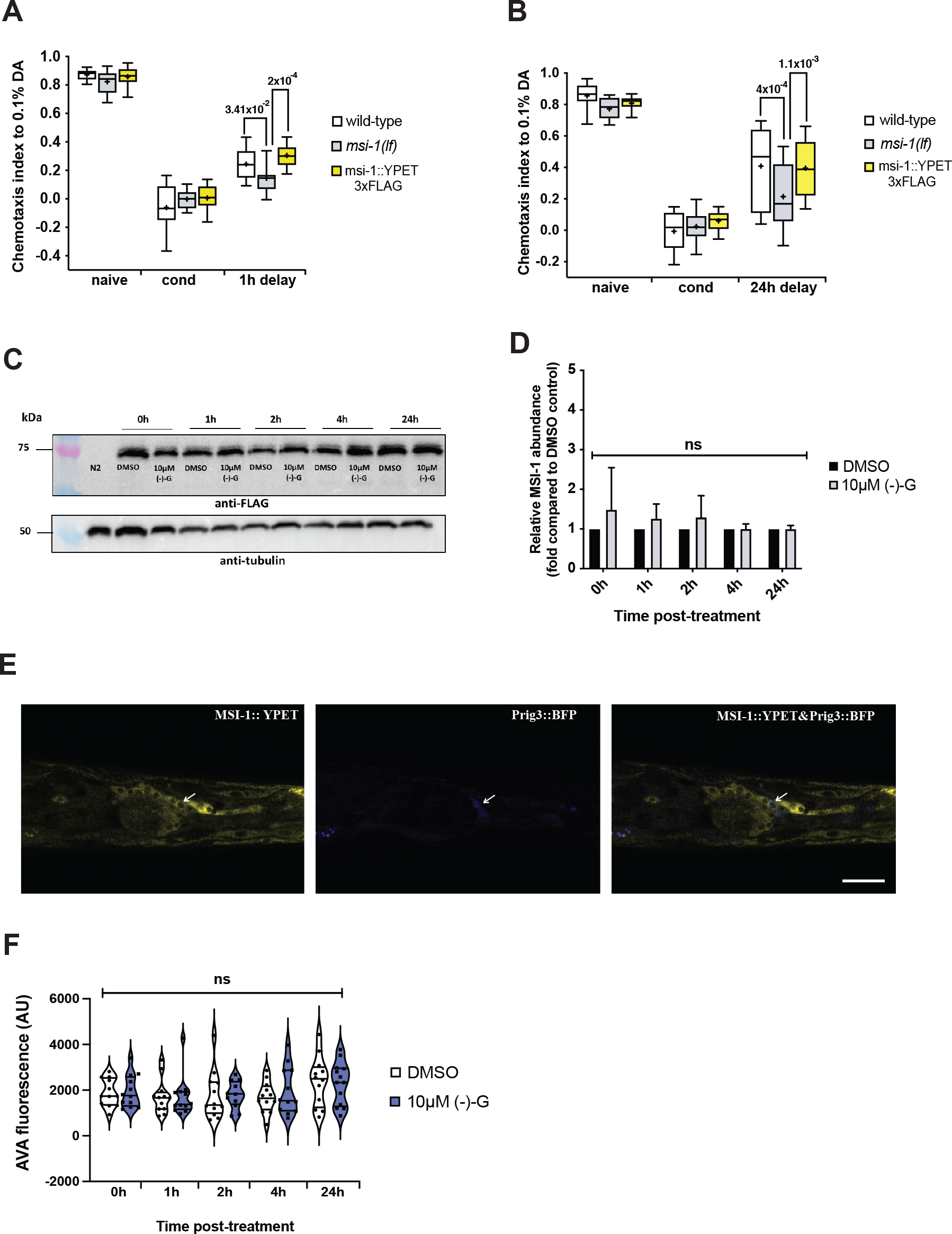
(-)- gossypol treatment does not influence MSI-1 protein abundance. (A) Negative STAM was tested in wild-type, *msi-1(lf)* and *msi-1∷3xFLAG∷YPET* worms. Worms were assayed toward DA without (naïve), with preincubation with DA and starvation (cond) or after 1h (1h delay). (B) Negative LTAM in the different genotypes was tested following two consecutive conditioning phases and DA preference was tested immediately (cond) and after 24h (24h delay) recovery period. All experiments were done in triplicates and repeated at least five times. Box plots for each genotype and condition are presented with whiskers indicating the 10th and 90th percentiles. Statistical significance between groups was assessed with two-way ANOVA and post hoc t-tests as indicated (Bonferroni’s adjusted p values are reported). (C) MSI-1 abundance following DMSO or (-)- gossypol treatment was analyzed in a time-dependent manner using western blots. For each condition, lysates from 50 transgenic worms were analyzed using FLAG antibody (top), and as the loading control membranes were reprobed for tubulin (bottom). (D) Relative MSI-1 abundance is plotted at different time points following a 2h -(−)gossypol treatment and values are presented as fold change to the DMSO control group. Bars represent mean ± SD from 6 independent blots. Statistical significance between groups was assessed with two-way ANOVA, ns: not significant. (E) Representative confocal images of MSI-1 expression in the AVA interneuron (arrows) in MSI-1∷YPET∷3xFLAG animals. Scale bar, 20μm. (F) Violin plots displaying quantification of fluorescence intensity in AVA interneuron in MSI-1∷YPET∷3xFLAG animals at different time points following DMSO and -(−)gossypol treatment. Animals were recorded with identical microscope settings and YPET intensity was measured on z-projected confocal images and quantified using ImageJ software. The centre line of the violin plot represents the group median, whereas the bottom and top lines represent the 25^th^ and 75^th^ percentiles respectively. Statistical analysis was performed using a two-way ANOVA, ns: not significant.

To check for possible (-)- gossypol-related changes in MSI-1 protein abundance, we treated worms for 2 hours with either DMSO or 10μM gossypol and collected worm fractions at different time points. By carrying out a western blot analysis, we could not detect any significant changes in MSI-1 protein levels (Figure 8C, D), suggesting that (-)- gossypol inhibits MSI-1 function without affecting its abundance. Since the fractions collected are whole worm lysates and represent MSI-1 global expression, we carried out fluorescent microscopy to identify any tissue-specific MSI-1 expression changes. We focused specifically on the AVA neuron and crossed the MSI-1∷YPET∷3xFLAG strain with an integrated *rig-3* promoter-driven BFP line to easily localize the AVA interneuron amongst the dense head neuronal network (Figure 8E). We could not detect any significant changes in MSI-1 expression at different time points following treatment, further corroborating our western blot results (Figure 8F).

### Effect of *msi-1* and (-)- gossypol on age-dependent cognitive decline

Physiological, age-dependent memory decline can be readily observed in nematodes with two-day-old worms already exhibiting a significant decline in olfactory associative memory (Kauffman et al. 2010). To investigate the potential beneficial effect of *msi-1* inhibition on age-dependent cognitive decline, we compared LTAM in one- and two-day-old wild-type and *msi-1 (lf)* worms. As expected, the LTAM of wild-type worms significantly decreased with age, whilst *msi-1 (lf)* displayed no age-dependent memory decline (Figure 9A). Furthermore, both one- and two-day-old *msi-1 (lf)* worms exhibited a significant increase in LTAM retention compared to their wild-type counterparts. In line with previous results, (-)- gossypol treatment of two-day-old wild-type worms mimicked the *msi-1 (lf)* phenotype, with animals displaying improved LTAM retention when compared to their DMSO treated counterparts (Figure 9B), suggesting that (-)- gossypol could help improve memory of aged worms.

**Figure 9.**
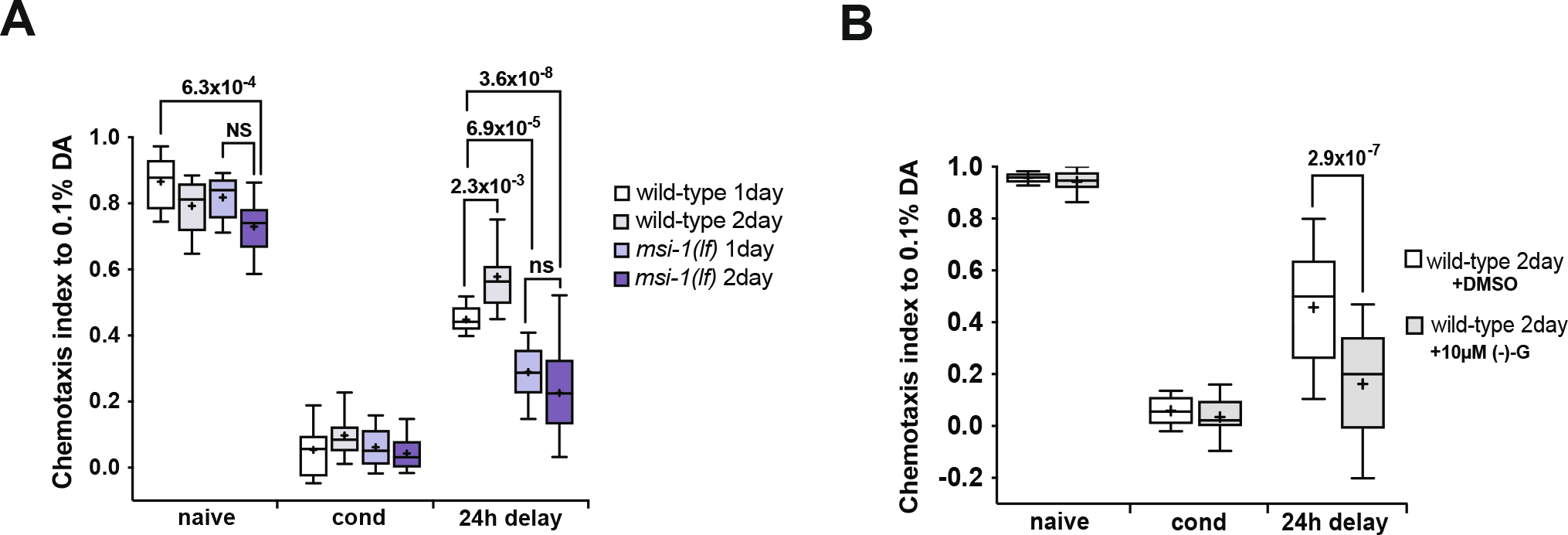
(-)- gossypol reverses physiological age-dependent cognitive decline in an MSI-1-dependent manner. (A) Negative LTAM was tested in 1-day and 2-day-old wild-type and *msi-1(lf)* worms and in (B) 2-day-old wild-type worms treated with DMSO and 10μM (-)- gossypol. Worms were assayed toward DA without (naïve), with two rounds of preincubation with DA and starvation (cond) or after 24h (24h delay). All experiments were done in triplicates and repeated at least five times. Box plots for each genotype and condition are presented with whiskers indicating the 10th and 90th percentiles. Statistical significance between groups was assessed with two-way ANOVA and post hoc t-tests as indicated (Bonferroni’s adjusted p values are reported).

## Discussion

A nervous system’s robust cognitive fitness does not stem solely from its ability to form new memories but also from its capacity to erase/forget irrelevant and harmful information (Davis and Zhong, 2017). Forgetting should not be regarded as having a negative connotation, but rather be seen as a dynamic cognitive process of the brain, enabling us to adapt and survive in a constantly changing environment. Thus, forgetting constitutes an innate system characterized by its own finely tuned molecular and cellular processes that should be distinguished from those regulating memory encoding, consolidation, and retrieval (Davis and Zhong, 2017). Here we report that the human musashi orthologues, MSI1 and MSI2, likely have a conserved role in the regulation of memory. Introducing human MSIs into *C. elegans* under the control of either the endogenous *msi-1* or *rig-3* promoter efficiently rescued the forgetting defect of *msi-1(lf)* worms, both in a short-term and long-term aversive associative learning paradigm. This suggests that despite the differences in the amino acid sequence between worm and human MSI proteins, their function seems to be conserved during the regulation of forgetting. This is the first study to suggest that human MSI proteins could have a role in the active regulation of forgetting. In addition, pharmacological treatment of human MSI1- and MSI2-expressing worms with the natural compound (-)- gossypol, improved memory with no effects on locomotion or memory acquisition of worms. MSI1 and MSI2 display a high degree of sequence and structure similarity and appear to target the same mRNA recognition motif (Ohyama et al., 2012). Therefore, (-)- gossypol could be considered as a MSI1/2 dual inhibitor.

The (-)- gossypol enantiomer used in this study is eliminated slowly (Yu-Wen, 1987; Wu et al., 1989), constitutes the most biologically active form of the compound, and consequently is considered more toxic than (+) gossypol (Bailey et al., 2000). Toxicity testing assays revealed no immediate or delayed toxic effects of (-)- gossypol concentrations up to 10μM, however, higher concentrations exhibited slight toxic effects, as illustrated by the reduced chemotactic response of worms. Additionally, (-)- gossypol did not alter the locomotory behavior of worms which could influence their response toward the chemoattractant diacetyl. However, in mammals, high concentrations of free gossypol have been shown to cause respiratory distress, impaired body weight gain, weakness, apathy, impaired reproduction, compromised immune function, and even death (Gadelha et al., 2014). Additionally, it should be pointed out, that the lack of toxicity in *C. elegans* does not preclude it in humans. Previous studies have shown how worm pharmacology can differ, where dopamine drugs had opposing effects on DA receptors in worms and mammals (Suo et al., 2004) and sodium azide is a permanent cytochrome c oxidase inhibitor in mammals whilst a reversible one in *C. elegans*. Thus, we acknowledge that caution must be paid when using *C. elegans* as a model for drug screening.

Furthermore, (-)- gossypol has been shown not to be selective against MSI but can act on multiple pathways (Cheng et al., 2003; Wang et al., 2013). However, we did not observe any behavioral changes in (-)- gossypol-treated *msi-1(lf)* worms. Moreover, (-)- gossypol treatment of worms expressing human MSI1 solely in the AVA interneuron inhibits the previously seen phenotypic rescue (Figure 3), suggesting a tissue-specific mode of drug action rather than an off-target effect. Nevertheless, while the concentrations used in our treatment might not suffice to affect other pathways, we cannot exclude that (-)- gossypol could harbor off-target effects, as increased toxicity is observed with increasing concentrations of the drug. Future efforts should, therefore, focus on elucidating compounds that selectively target and inhibit MSI1/2.

Finally, physiological age-dependent cognitive decline normally observed in wild-type worms did not occur in two-day-old *msi-1(lf)* worms or two-day-old wild-type worms treated with (-)- gossypol. These results implicate musashi in age-dependent memory decline and nominate musashi inhibiting compounds as memory modulators.

Taken together, our results argue in favor of an evolutionary conserved role for musashi proteins in memory in species as diverse as nematodes and humans. Additionally, we show that *C. elegans,* similar to other model organisms (Papanikolopoulou et al., 2019), can be successfully used as a model for therapeutic intervention using pharmacological manipulations (Giunti et al., 2021), which might prove useful in the treatment of memory-related disorders.

## Methods

### General methods and strains used

Common reagents were obtained from Sigma (Sigma-Aldrich, St Louis, MO) unless otherwise indicated. Standard methods were used for maintaining and manipulating *C. elegans* (Brenner, 1974). All experiments were conducted using a synchronized population of young adult hermaphrodites. The *C. elegans* Bristol strain, variety N2, was used as the wild-type reference strain in all experiments. Alleles and transgenes used were: *msi-1(os1), msi-1(os1);utrIs15[p_msi-1_∷human msi1cDNA∷msi-1 3’UTR, p_sur-5_∷mDsred], msi-1(os1);utrIs14[p_msi-_1∷human msi2cDNA∷msi1 3’UTR, p_sur-5_∷mDsred], msi-1(os1);utrIs40[p_rig-3_∷human msi1cDNA∷msi-1 3’UTR, p_sur-5_∷mDsred], msi-1 (utr55[YPET∷3xFLAG]), utrSi43[prig-3∷LoxP∷BFP∷LoxP∷FLP-DS∷SL2∷GFP∷H2B],unc-119(ed3), msi-1(utr55[YPET∷3xFLAG]);utrSi43[prig-3∷LoxP∷BFP∷LoxP∷FLP-DS∷SL2∷GFP∷H2B],unc-119(ed3), utrIs11[p_msi-1_∷GFP∷arx-23’UTR, punc-119+].*

Extrachromosomal transgenic lines were generated by injecting DNA at a concentration of 100 ng/μl into both arms of the syncytial gonad of worms as previously described (Mello et al., 1991). *psur-5∷mDsRed* was used as a transformation marker at a concentration of 10 ng/μl. Chromosomal integration of extrachromosomal arrays was done by UV radiation for 12s or 15s. Following integration, generated strains were backcrossed four times to the wild-type strain.

### *C. elegans* behavior assays

Chemotaxis to different compounds was assessed in synchronized young adult worms as described previously (Bargmann et al, 1993). Briefly, a population of well-fed, young adults was washed three times with CTX buffer (5 mM KH_2_PO_4_/K_2_HPO_4_ pH 6.0, 1 mM CaCl_2_, and 1 mM MgSO_4_) and 100–200 worms were placed in the center of a 10-cm CTX test plate. Worms were given a choice between a spot of attractant diluted in ethanol with 20 mM sodium-azide and a counter spot with ethanol and sodium-azide (NaN_3_). The distribution of the worms over the plate was determined after 1h and the chemotaxis index (CI) was calculated as previously described 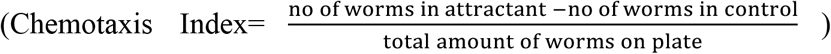 (Bargmann et al, 1993). Olfactory conditioning was assessed as described (Nuttley et al. 2002), with a few modifications. Starvation conditioning was performed without food in the presence of 2 μl undiluted diacetyl (DA) spotted on the plate for 1 h on 10 cm CTX plates (5 mM KH_2_PO_4_/K_2_HPO_4_ pH 6.0, 1 mM CaCl_2_, 1 mM MgSO_4_, 2% agar). Following conditioning, worms were tested for their chemotaxis towards DA immediately after or after 1h rest to assess short-term associative memory.

Long-term associative memory was induced with two cycles of conditioning with DA with 30 minutes of rest between trainings. Following conditioning, worms were kept on NGM plates in the presence of abundant food for 24 h and tested for chemotaxis to DA after the recovery phase (Vukojevic et al., 2012).

Pharmacological treatment was performed by soaking worms for 2 hours in CTX supplemented with either dimethyl sulfoxide (DMSO) alone or the indicated concentration of gossypol (AT101, Sigma-Aldrich) in DMSO, prior to olfactory conditioning. Gossypol was dissolved in absolute DMSO to make a 10mM stock solution. After a 2h incubation with the drug, worms were washed 3 times with CTX and tested for their preference to DA immediately (naïve), after conditioning, 1h (STAM), or 24h (LTAM) after conditioning. Furthermore, any potential delayed drug toxicity was tested 2h and 24h after treatment with the drug by assessing the chemotaxis to DA.

### Locomotory Rate Assays

Locomotory rate assays were carried out to test for any gossypol-related effects on worm motility. After treatment, worms were allowed to recover for 30 minutes on plates seeded with OP50. Individual worms were transferred on 6 cm assay plates, allowed to move freely for 3 minutes and then the number of body bends of fifteen animals from each strain and treatment group as indicated was counted for 1 minute.

### Molecular biology

Rescue of the *msi-1(lf)* phenotype was performed with a 12.9 kb construct comprised of a 7.7 kb fragment of the *C. elegans msi-1* promoter region, the full-length *msi1* or *msi2* human cDNA designed with NcoI and KpnI compatible ends, fused with a 1.1 kb fragment of the *C. elegans msi-1* 3’UTR. Full-length human *msi1* and *msi2* cDNA were amplified from the MGC clones pENTR™223.1 and pOTB7 respectively (Transomic Technologies, Huntsville, AL United States). Primers used for human *msi1* cDNA amplification were: CCA TGG AGA CTG ACG CGC CCC AG and GGT ACC TCA GTG GTA CCC ATT GGT GAA GG and for human *msi2* cDNA amplification CCA TGG AGG CAA ATG GGA GCC AAG and GGT ACC TCA ATG GTA TCC ATT TGT AAA GG. The thermal cycle was programmed for 120 seconds at 95°C as initial denaturation, followed by 30 cycles of 30s at 95°C for denaturation, 30s at 64 °C as annealing, 90s at 72 °C for extension, and final extension at 72 °C for 5 min. For the tissue-specific rescue experiments, the full-length *msi1* or *msi2* human cDNA and the *msi-1* 3’UTR were fused to a 3 kb fragment of the *rig-3* promoter using NEBuilder Hifi DNA assembly (New England Biolabs, Ipswich, MA).

### Reverse Transcription PCR (RT-PCR)

Total RNA was isolated from synchronized adult wild-type, *hmsi1* and *hmsi2* transgene-carrying animals using a Direct-zol RNA MiniPrep kit (Zymo Research Cooperation) with DNase treatment. RNA integrity was measured using the Agilent 2100 Bioanalyzer (Agilent Genomics, California, United States). cDNA was synthesized using SuperScript™ III First-Strand Synthesis System (ThermoFischer Scientific, MA, USA) according to the manufacturer’s recommendations using 1μg of purified RNA. Primers designed to amplify human musashi cDNA were the following: for *msi-1* ATT GAC CCT AAG GTG GCC TTC C and TGG CGG CGC TGA TGT AAC TG and for *msi-2* ACG TTC GCA GAC CCA GCA AG and TGG GAA GCC TGG GAA CTG ATA G. Plasmid DNA was used to test for primer annealing. To avoid amplification from contaminating genomic DNA, two sets of primers per construct were designed to span the endogenous *C. elegans msi1* promoter and human cDNA boundary region (*hmsi1*: ACG AAA CAA ACC ATG GAG AC and TGC GTA GTC TGC CAA CTG AG; CAA ACC ATG GAG ACT GAC GC and GAA GTA TTC GCG CAG CCC TT *hmsi2*: ACG AAA CAA ACC ATG GAG GCA and GGC TAT CTG GTG AGG TCT GC; CAA ACC ATG GAG GCA AAT GGG and CTA AGG CTA TCT GGT GAG GTC. Actin was used as a control for cDNA amplification with the following primers (forward primer: GCC CAA TCC AAG AGA GGT ATC C, reverse primer: GGC AAC ACG AAG CTC ATT G). The thermal cycle was programmed for 120 seconds at 95°C as initial denaturation, followed by 30 cycles of 30s at 95°C for denaturation, 30s at 60 °C as annealing, 60s at 72 °C for extension, and final extension at 72 °C for 5 min.

### Targeted modification of *msi-1* using CRISPR/Cas9

Endogenous tagging of *msi-1* with YPET∷3xFLAG was generated as described previously (Dickinson et al., 2015). 601bp and 722bp homology arms flanking the N-terminus of *msi-1* were PCR amplified from N2 genomic DNA and inserted into the mNG^SEC&3xFLAG vector pDD283 using NEBuilder Hifi DNA assembly (New England Biolabs, Ipswich, MA). The Cas9 target site was selected using the Sequence Scan for CRISPR database (http://cistrome.org/SSC/) and inserted into pDD162 (Dickinson et al, 2013). The sgRNA sequence used was 5’-ATGACAACGACAGTATCAACGTTTTAGAGCTAGAAATAGCAAGT-3’. A mixture of 50 ng/μl Cas9–sgRNA plasmid, 10 ng/μl repair template, and 2.5 ng/μl pCFJ90, 5 ng/μl pCFJ104, and 10 ng/μl *sur-5p∷dsRed* co-injection markers were injected into the gonads of young adults (Mello et al., 1991). The knock-in line was established, the SEC cassette was excised using heat shock and the *msi-1* YPET∷3xFLAG line was sequenced to verify correct insertion of the tag.

### Western blot analysis

For protein extraction, 50 worms per condition were collected in ice-cold 3×Laemmli buffer (6%w/v SDS, 30%glycerol, 120mM Tris-Cl pH 6.8, 0.03% w/v bromophenol blue) and heated for 5 mins at 95°C. Total lysate samples were subjected to SDS-PAGE, transferred to PVDF membranes, blocked with 5% nonfat dry milk in TBST (50mM Tris-HCl, pH 7.5, 150mM NaCl, 0.05% Tween-20), and incubated with primary antibodies as indicated. Antibodies used were: mouse anti-FLAG (1:2000, Sigma Aldrich, St. Louis, MI) and mouse anti-tubulin (1:20000, Merck Millipore, Burlington, MA). Primary antibodies were detected using HRP coupled secondary antibodies (1:10000, Jackson ImmunoResearch Laboratories, Cambridge House, UK). Chemiluminescent signal was developed using Clarity and ClarityMax Western Blotting Substrates (BioRad Laboratories Inc., Hercules, CA) followed by detection with a FujiFilm ImageQuant LAS-4000 detector (GE Healthcare, Chicago, IL).

### Fluorescence Microscopy

Whole worms were mounted on 3% agarose pads and immobilized with 0.5% NaN_3_. Synchronized one-day-old adult worms were imaged using a Zeiss LSM 880 inverted scanning confocal microscope equipped with 25× and 63× oil immersion objectives. All recorded images were processed and quantified using Fiji software (version 2.1.0/1.53c). Fluorescence in AVA was measured using the integrated density function and the corresponding background signal was subtracted. Due to the dense neuronal signal in the head of the animal, AVA was first localized and selected in the BFP channel and the identical selection was then superimposed onto the YPET channel.

### Statistics

All data and statistical analyses were carried out using the Prism software (version: 9.1.2). . Main effects and interaction terms were tested using ANOVA. Statistical tests for significance were done with F-tests using sum-of-squares type I. The p-value threshold was set to nominal significance (p < 0.05). In case of a significant main or interaction effect, significance between data was tested using post-hoc t-tests. P-values of the post-hoc tests were corrected for the number of tests calculated per analysis (Bonferroni-correction per analysis: p_bonf_ < 0.05). For the imaging quantification, a two-tailed unpaired Student’s t-test was carried out to assess any differences between groups, and the p-value threshold was set to nominal significance (p < 0.05).

## Acknowledgments

We are grateful to Anne Spang for generously sharing reagents and instruments. We would also like to thank the Caenorhabditis Genetic Center (supported by NIH-NCRR) for providing nematode strains. The work was supported by the Swiss National Science Foundation (SNSF) grant (31003A_156579). We would finally like to thank the Imaging Core Facility of the Biozentrum (University of Basel) for microscopy support.

## Author contributions

PM designed and performed the experiments, evaluated the results, and wrote the manuscript; AA designed and performed the experiments; FP performed the experiments; CS, DQ, AS, and AP evaluated the results and wrote the manuscript.

## References

1. Bailey, C. A., Stipanovic, R. D., Ziehr, M. S., Haq, A. U., Sattar, M., Kubena, L. F., Kim, H. L., & de M. Vieira, R. (2000). Cottonseed with a High (+)- to (−)-Gossypol Enantiomer Ratio Favorable to Broiler Production. Journal of Agricultural and Food Chemistry, 48(11), 5692–5695. https://doi.org/10.1021/jf000211n

2. Bargmann, C. I., Hartwieg, E., & Horvitz, H. R. (1993). Odorant-selective genes and neurons mediate olfaction in C. elegans. Cell, 74(3), 515–527. https://doi.org/10.1016/0092-8674(93)80053-H

3. Bennett, C. G., Riemondy, K., Chapnick, D. A., Bunker, E., Liu, X., Kuersten, S., & Yi, R. (2016). Genome-wide analysis of Musashi-2 targets reveals novel functions in governing epithelial cell migration. Nucleic Acids Research, 44(8), 3788–3800. https://doi.org/10.1093/nar/gkw207

4. Brenner, S. (1974). THE GENETICS OF *CAENORHABDITIS ELEGANS*. Genetics, 77(1), 71–94. https://doi.org/10.1093/genetics/77.1.71

5. Cheng, J.-S., Lo, Y.-K., Yeh, J.-H., Cheng, H.-H., Liu, C.-P., Chen, W.-C., & Jan, C.-R. (2003). Effect of gossypol on intracellular Ca2+ regulation in human hepatoma cells. The Chinese Journal of Physiology, 46(3), 117–122.

6. Clingman, C. C., Deveau, L. M., Hay, S. A., Genga, R. M., Shandilya, S. M., Massi, F., & Ryder, S. P. (2014). Allosteric inhibition of a stem cell RNA-binding protein by an intermediary metabolite. ELife, 3, e02848. https://doi.org/10.7554/eLife.02848

7. Colbert, H. A., & Bargmann, C. I. (1995). Odorant-specific adaptation pathways generate olfactory plasticity in C. elegans. Neuron, 14(4), 803–812. https://doi.org/10.1016/0896-6273(95)90224-4

8. Colitti, M., & Farinacci, M. (2009). Expression of a putative stem cell marker, Musashi 1, in mammary glands of ewes. Journal of Molecular Histology, 40(2), 139–149. https://doi.org/10.1007/s10735-009-9224-3

9. Darnell, R. B. (2010). HITS-CLIP: Panoramic views of protein–RNA regulation in living cells. Wiley Interdisciplinary Reviews: RNA, 1(2), 266–286. https://doi.org/10.1002/wrna.31

10. Davis, R. L., & Zhong, Y. (2017). The Biology of Forgetting—A Perspective. Neuron, 95(3), 490–503. https://doi.org/10.1016/j.neuron.2017.05.039

11. Dickinson, D. J., Ward, J. D., Reiner, D. J., & Goldstein, B. (2013). Engineering the Caenorhabditis elegans genome using Cas9-triggered homologous recombination. Nature Methods, 10(10), 1028–1034. https://doi.org/10.1038/nmeth.2641

12. Dickinson, D. J., Pani, A. M., Heppert, J. K., Higgins, C. D., & Goldstein, B. (2015). Streamlined Genome Engineering with a Self-Excising Drug Selection Cassette. Genetics, 200(4), 1035–1049. https://doi.org/10.1534/genetics.115.178335

13. Gadelha, I. C. N., Fonseca, N. B. S., Oloris, S. C. S., Melo, M. M., & Soto-Blanco, B. (2014). Gossypol Toxicity from Cottonseed Products. The Scientific World Journal, 2014, 1–11. https://doi.org/10.1155/2014/231635

14. Giunti, S., Andersen, N., Rayes, D., & De Rosa, M. J. (2021). Drug discovery: Insights from the invertebrate *Caenorhabditis elegans*. Pharmacology Research & Perspectives, 9 (2). https://doi.org/10.1002/prp2.721

15. Hadziselimovic, N., Vukojevic, V., Peter, F., Milnik, A., Fastenrath, M., Fenyves, B. G., Hieber, P., Demougin, P., Vogler, C., de Quervain, D. J.-F., Papassotiropoulos, A., & Stetak, A. (2014). Forgetting Is Regulated via Musashi-Mediated Translational Control of the Arp2/3 Complex. Cell, 156(6), 1153–1166. https://doi.org/10.1016/j.cell.2014.01.054

16. Kandel, E. R., Dudai, Y., & Mayford, M. R. (2014). The Molecular and Systems Biology of Memory. Cell, 157(1), 163–186. https://doi.org/10.1016/j.cell.2014.03.001

17. Karmakar, S., Ramirez, O., Paul, K. V., Gupta, A. K., Kumari, V., Botti, V., de los Mozos, I. R., Neuenkirchen, N., Ross, R. J., Karanicolas, J., Neugebauer, K. M., & Pillai, M. M. (2022). Integrative genome-wide analysis reveals EIF3A as a key downstream regulator of translational repressor protein Musashi 2 (MSI2). NAR Cancer, 4 (2), zcac015. https://doi.org/10.1093/narcan/zcac015

18. Kauffman, A. L., Ashraf, J. M., Corces-Zimmerman, M. R., Landis, J. N., & Murphy, C. T. (2010). Insulin Signaling and Dietary Restriction Differentially Influence the Decline of Learning and Memory with Age. PLoS Biology, 8 (5), e1000372. https://doi.org/10.1371/journal.pbio.1000372

19. Kayahara, T., Sawada, M., Takaishi, S., Fukui, H., Seno, H., Fukuzawa, H., Suzuki, K., Hiai, H., Kageyama, R., Okano, H., & Chiba, T. (2003). Candidate markers for stem and early progenitor cells, Musashi-1 and Hes1, are expressed in crypt base columnar cells of mouse small intestine. FEBS Letters, 535 (1–3), 131–135. https://doi.org/10.1016/S0014-5793(02)03896-6

20. Kharas, M. G., Lengner, C. J., Al-Shahrour, F., Bullinger, L., Ball, B., Zaidi, S., Morgan, K., Tam, W., Paktinat, M., Okabe, R., Gozo, M., Einhorn, W., Lane, S. W., Scholl, C., Fröhling, S., Fleming, M., Ebert, B. L., Gilliland, D. G., Jaenisch, R., & Daley, G. Q. (2010). Musashi-2 regulates normal hematopoiesis and promotes aggressive myeloid leukemia. Nature Medicine, 16(8), 903–908. https://doi.org/10.1038/nm.2187

21. Kraemer, P. J., & Golding, J. M. (1997). Adaptive forgetting in animals. Psychonomic Bulletin & Review, 4(4), 480–491. https://doi.org/10.3758/BF03214337

22. Lan, L., Appelman, C., Smith, A. R., Yu, J., Larsen, S., Marquez, R. T., Liu, H., Wu, X., Gao, P., Roy, A., Anbanandam, A., Gowthaman, R., Karanicolas, J., De Guzman, R. N., Rogers, S., Aubé, J., Ji, M., Cohen, R. S., Neufeld, K. L., & Xu, L. (2015). Natural product (−)-gossypol inhibits colon cancer cell growth by targeting RNA-binding protein Musashi-1. Molecular Oncology, 9(7), 1406–1420. https://doi.org/10.1016/j.molonc.2015.03.014

23. Lan, L., Liu, H., Smith, A. R., Appelman, C., Yu, J., Larsen, S., Marquez, R. T., Wu, X., Liu, F. Y., Gao, P., Gowthaman, R., Karanicolas, J., De Guzman, R. N., Rogers, S., Aubé, J., Neufeld, K. L., & Xu, L. (2018). Natural product derivative Gossypolone inhibits Musashi family of RNA-binding proteins. BMC Cancer, 18 (1), 809. https://doi.org/10.1186/s12885-018-4704-z

24. Lian, J., Ni, Z., Dai, X., Su, C., Smith, A. R., Xu, L., & He, F. (2012). Sorafenib Sensitizes (−)-Gossypol-Induced Growth Suppression in Androgen-Independent Prostate Cancer Cells via Mcl-1 Inhibition and Bak Activation. Molecular Cancer Therapeutics, 11(2), 416–426. https://doi.org/10.1158/1535-7163.MCT-11-0559

25. Lukong, K. E., Chang, K., Khandjian, E. W., & Richard, S. (2008a). RNA-binding proteins in human genetic disease. Trends in Genetics, 24(8), 416–425. https://doi.org/10.1016/j.tig.2008.05.004

26. Mello, C. C., Kramer, J. M., Stinchcomb, D., & Ambros, V. (1991). Efficient gene transfer in C.elegans: Extrachromosomal maintenance and integration of transforming sequences. The EMBO Journal, 10(12), 3959–3970.

27. Nuttley, W. M., Atkinson-Leadbeater, K. P., & van der Kooy, D. (2002). Serotonin mediates food-odor associative learning in the nematode Caenorhabditis elegans. Proceedings of the National Academy of Sciences, 99(19), 12449–12454. https://doi.org/10.1073/pnas.192101699

28. Ohyama, T., Nagata, T., Tsuda, K., Kobayashi, N., Imai, T., Okano, H., Yamazaki, T., & Katahira, M. (2012). Structure of Musashi1 in a complex with target RNA: The role of aromatic stacking interactions. Nucleic Acids Research, 40(7), 3218–3231. https://doi.org/10.1093/nar/gkr1139

29. Okano, H., Kawahara, H., Toriya, M., Nakao, K., Shibata, S., & Imai, T. (2005). Function of RNA-binding protein Musashi-1 in stem cells. Experimental Cell Research, 306(2), 349–356. https://doi.org/10.1016/j.yexcr.2005.02.021

30. Oliver, C. L. (2004). In vitro Effects of the BH3 Mimetic, (−)-Gossypol, on Head and Neck Squamous Cell Carcinoma Cells. Clinical Cancer Research, 10(22), 7757–7763. https://doi.org/10.1158/1078-0432.CCR-04-0551

31. Papanikolopoulou, K., Mudher, A., & Skoulakis, E. (2019). An assessment of the translational relevance of Drosophila in drug discovery. Expert opinion on drug discovery, 14(3), 303–313. https://doi.org/10.1080/17460441.2019.1569624

32. Ryan, T. J., & Frankland, P. W. (2022). Forgetting as a form of adaptive engram cell plasticity. Nature reviews. Neuroscience, 23(3), 173–186. https://doi.org/10.1038/s41583-021-00548-3

33. Sakakibara, S., Imai, T., Hamaguchi, K., Okabe, M., Aruga, J., Nakajima, K., Yasutomi, D., Nagata, T., Kurihara, Y., Uesugi, S., Miyata, T., Ogawa, M., Mikoshiba, K., & Okano, H. (1996). Mouse-Musashi-1, a Neural RNA-Binding Protein Highly Enriched in the Mammalian CNS Stem Cell. Developmental Biology, 176(2), 230–242. https://doi.org/10.1006/dbio.1996.0130

34. Sengupta, P., Chou, J. H., & Bargmann, C. I. (1996). odr-10 encodes a seven transmembrane domain olfactory receptor required for responses to the odorant diacetyl. Cell, 84(6), 899–909. https://doi.org/10.1016/s0092-8674(00)81068-5

35. Squire, L. R. (1987). The organization and neural substrates of human memory. International Journal of Neurology, 21–22, 218–222.

36. Suo, S., Ishiura, S., & Van Tol, H. H. M. (2004). Dopamine receptors in C. elegans. European Journal of Pharmacology, 500 (1–3), 159–166. https://doi.org/10.1016/j.ejphar.2004.07.021

37. Vukojevic, V., Gschwind, L., Vogler, C., Demougin, P., de Quervain, D. J.-F., Papassotiropoulos, A., & Stetak, A. (2012). A role for α-adducin (ADD-1) in nematode and human memory: α-Adducin regulates synaptic plasticity. The EMBO Journal, 31(6), 1453–1466. https://doi.org/10.1038/emboj.2012.14

38. Wang, J., Jin, L., Li, X., Deng, H., Chen, Y., Lian, Q., Ge, R., & Deng, H. (2013). Gossypol induces apoptosis in ovarian cancer cells through oxidative stress. Molecular BioSystems, 9 (6), 1489. https://doi.org/10.1039/c3mb25461e

39. Wang, S.-H., & Morris, R. G. M. (2010). Hippocampal-Neocortical Interactions in Memory Formation, Consolidation, and Reconsolidation. Annual Review of Psychology, 61(1), 49–79. https://doi.org/10.1146/annurev.psych.093008.100523

40. Wu, D. (1989). An Overview of the Clinical Pharmacology and Therapeutic Potential of Gossypol as a Male Contraceptive Agent and in Gynaecological Disease: Drugs, 38(3), 333–341. https://doi.org/10.2165/00003495-198938030-00001

41. Yang, Z., Lan, L., Wu, X., Xu, L., & Buechner, M. (2019). RNA-Binding Proteins MSI-1 (Musashi) and EXC-7 (HuR) Regulate Serotonin-Mediated Behaviors in *C. elegans*. BioRxiv, 748509. https://doi.org/10.1101/748509

42. Yoda, A., Sawa, H., & Okano, H. (2000). MSI-1, a neural RNA-binding protein, is involved in male mating behavior in Caenorhabditis elegans. Genes to Cells, 5(11), 885–895. https://doi.org/10.1046/j.1365-2443.2000.00378.x

43. Yu-Wen, Y. (1987). Probing into the mechanism of action, metabolism, and toxicity of gossypol by studying its (+)- and (−)-stereoisomers. Journal of Ethnopharmacology, 20(1), 65–78. https://doi.org/10.1016/0378-8741(87)90120-6

44. Zearfoss, N. R., Deveau, L. M., Clingman, C. C., Schmidt, E., Johnson, E. S., Massi, F., & Ryder, S. P. (2014). A Conserved Three-nucleotide Core Motif Defines Musashi RNA Binding Specificity. Journal of Biological Chemistry, 289(51), 35530–35541. https://doi.org/10.1074/jbc.M114.597112

